# Viscoelastic HyA Hydrogel Promotes Recovery of Muscle Quality and Vascularization in a Murine Model of Delayed Rotator Cuff Repair

**DOI:** 10.1101/2025.01.08.632047

**Authors:** Morgan R. Pfaff, Aboubacar Wague, Michael Davies, Anouk R. Killaars, Derek Ning, Steven Garcia, Anthony Nguyen, Prashant Nuthalapati, Mengyao Liu, Xuhui Liu, Brian T. Feeley, Kevin E. Healy

## Abstract

Rotator cuff tears are among the most common musculotendinous injuries with high risk of permanent functional disability. Following surgical repair, sub-optimal patient outcomes are directly correlated with poor muscle quality; namely, injury site fatty infiltration (FI), fibrosis, and muscle atrophy. Muscle resident fibro-adipogenic progenitor cells (FAPs) have been identified as key regulators of post-injury skeletal muscle regeneration and repair by maintaining a promyogenic environment. In this work, human-derived FAPs (hFAPs) were encapsulated into hyaluronic acid (HyA)-based hydrogels functionalized with bsp-RGD(15) cell adhesion peptide, heparin, and a matrix metalloproteinase (MMP)-cleavable crosslinker. Hydrogel-encapsulated hFAPs increased expression of the promyogenic marker UCP1 and production of the anti-inflammatory cytokine IL-10, while downregulating the expression of the fibrotic marker αSMA over time. A murine model of unilateral rotator cuff transection, denervation, and delayed repair was treated with the HyA hydrogel or PBS and compared to a contralateral, non-injured control limb. Muscle histology 6 weeks post-repair revealed that the hydrogel reduced fibrosis, FI, and muscle atrophy while supporting vascularization of the injured tissue region. Collectively, these results suggest that the hydrogel alone can promote muscle regeneration in a clinically relevant delayed repair model of rotator cuff tear, which we hypothesize is due to controlled FAP differentiation into promyogenic lineages.

## 1. Introduction

Rotator cuff injuries are one of the most common upper extremity causes of physician visits in the United States, ranking only behind back and neck pain. Rotator cuff tears cause pain and loss of function, and their incidence is continuously rising in an aging population. Indeed, up to 20% of patients over the age of 50 show evidence of a full-thickness rotator cuff tear; by the age of 70, this prevalence jumps to 49%.^[1]^ Several studies have demonstrated that injury site muscle quality largely governs post-repair outcomes.^[2,3]^ Muscle atrophy and secondary muscle degeneration indicators including fibrosis and fatty infiltration (FI) are directly correlated with higher risk of re-tear and decreased functional recovery.^[4–6]^ With approximately 25-35% of patients exhibiting clinically significant amounts of atrophy and FI,^[7,8]^ developing treatment strategies to improve muscle quality and post-repair outcomes represents a promising avenue to alleviate an ever-increasing clinical burden.

Prior studies have identified fibro-adipogenic progenitor cells (FAPs), non-myogenic muscle resident stromal cells, as key regulators in repair of injured skeletal muscles,^[9,10]^ making them powerful targets for biomaterial therapeutic development. Upon injury, quiescent FAPs activate and rapidly expand in response to pro-inflammatory signals. The multipotent nature of FAPs drives their dynamic control over the regenerative niche (**Figure 1**). FAPs drive post-injury fibrosis and FI by differentiating into fibroblasts and white adipose tissue (WAT) cells, respectively. Excessive extracellular matrix (ECM) deposition by fibroblasts can hinder muscle function by reducing contractile tissue area and increase the risk of re-injury,^[11]^ while WAT accumulation is linked to myopathy and may influence the muscle environment as an endocrine regulator.^[12]^ Under β-adrenergic signaling, FAPs can adopt a promyogenic brown adipose tissue (BAT) phenotype known as brown FAPs (bFAPs).^[13]^ bFAPs have been shown to secrete promyogenic growth factors and exosomes, providing muscle satellite cells (MuSCs) with signals to expand, differentiate, and fuse into myotubes while coordinating inflammatory responses to injury.^[14]^ It has been previously demonstrated that transplantation of bFAPs into an *in vivo* rotator cuff injury model significantly improved muscle quality and function while reducing degeneration.^[15]^ Therefore, a therapeutic capable of inducing BAT differentiation of FAPs while minimizing the anti-myogenic effects of repair processes like fibrosis and FI would be of great interest for implementation as rotator cuff tear treatments.

**Figure 1:**
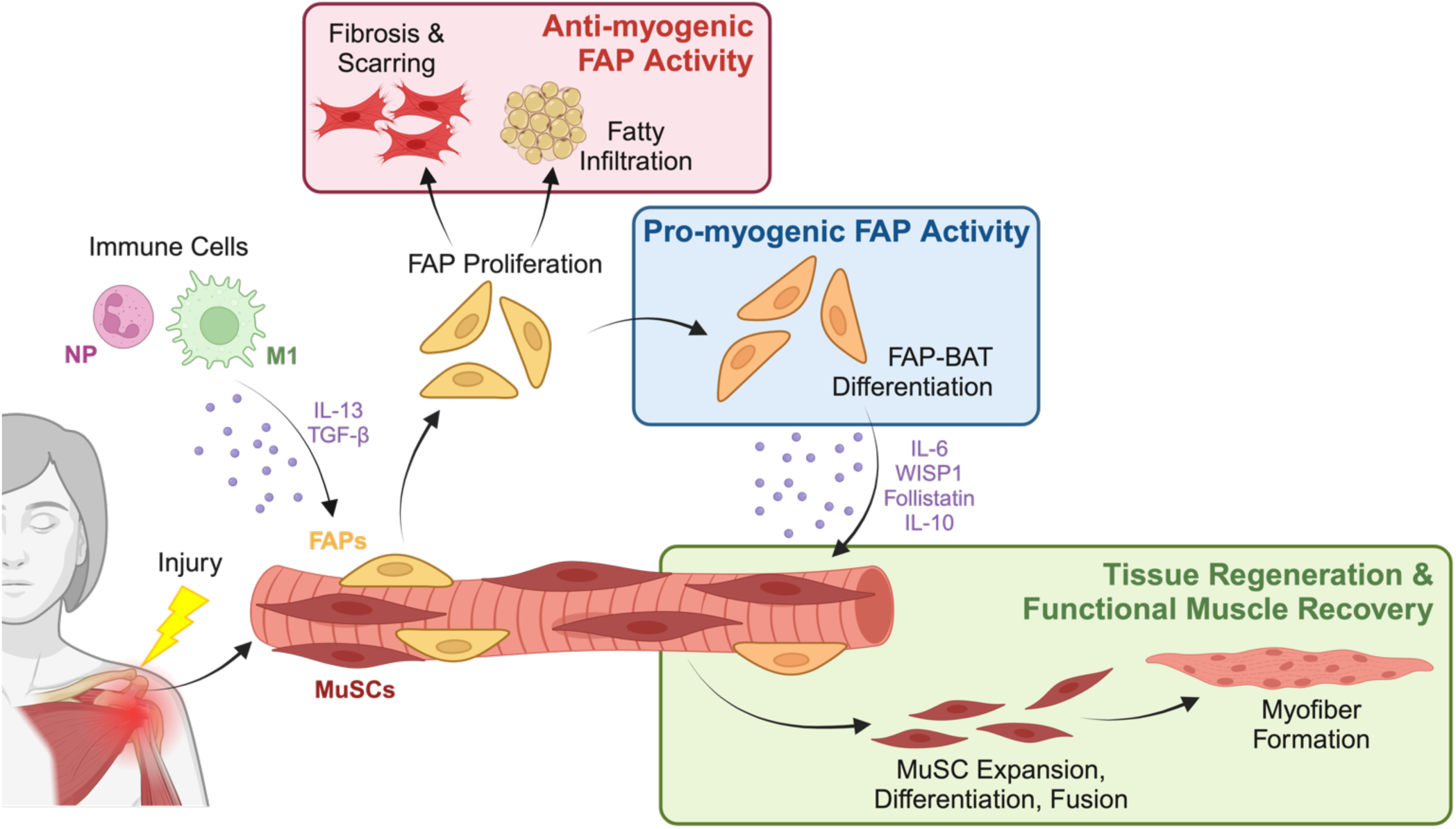
Regenerative and Degenerative Roles of FAPs in Muscle Regeneration. Upon injury, immune cells signal for muscle resident FAPs to expand. bFAP differentiation leads promyogenic activity such as secretion of factors that aid in muscle regeneration by signaling for satellite cell expansion, differentiation, and fusion into myotubes. Simultaneously, FAPs can also differentiate into fibroblasts and white fat that form non-contractile scar tissue and white fat, respectfully, thereby hindering regeneration. NP = neutrophil, M1 = M1 macrophage, FAP = fibro-adipogenic progenitor cells, MuSCs = muscle satellite cells.

Substantial efforts have been made to create biomaterial scaffolds for rotator cuff repair that promote multi-tissue level healing and regeneration.^[16,17]^ Frequently used as delivery systems for drugs,^[18]^ growth factors,^[19,20]^ or cells,^[21,22]^ biomaterial-based therapeutics aim to provide promyogenic cues and ECM-like mechanical support while limiting undesirable FI and scar tissue formation. Biocompatible, natural polymers such as hyaluronic acid (HyA) combined with engineering approaches to incorporate cell specificity, growth factor retention, and controlled degradation have the potential to recapitulate the muscle microenvironment and provide the appropriate, timely cues to direct biological activity. In addition to its ease of chemical modification, HyA is a major component of native ECM and plays crucial roles in native tissue repair and development.^[23,24]^ The body’s naturally-produced hyaluronidase enzymes enable complete biodegradation of HyA-based biomaterials as it is eventually replaced with functional tissue, making HyA an attractive platform to build promyogenic devices.

To this end, recent work in our laboratory has focused on the development of a semi-synthetic, modular HyA hydrogel which has demonstrated remarkable pro-regenerative behavior both *in vitro* and *in vivo*. The HyA polymer backbone is decorated with cell adhesion peptides for cell specificity and mobility, as well as high molecular weight heparin to sequester growth factors released by infiltrating cells. Crosslinking of the system via a matrix metalloproteinase (MMP)-cleavable, dithiol crosslinker fosters cell-mediated matrix remodeling and degradation tailored to the rate of cell infiltration and ECM deposition (**Figure 2A**). Scaffold components are bound to the acrylated HyA (acHyA) backbone via Michael-type addition, enabled by thiolation of heparin and addition of terminal, thiol-containing cysteines to adhesion and MMP-cleavable peptides. In addition to promoting the engraftment of transplanted stem cells in a murine model, this HyA hydrogel system has demonstrated robust angiogenic behavior *in vitro* and the ability to promote vascularization *in vivo*.^[25–30]^ Furthermore, significant functional recovery was observed using a rat tibialis anterior volumetric muscle loss model implanted with this hydrogel, as evaluated by nerve-stimulated muscle contraction and histological assessment of the muscle 12 weeks post-repair.^[30]^

**Figure 2:**
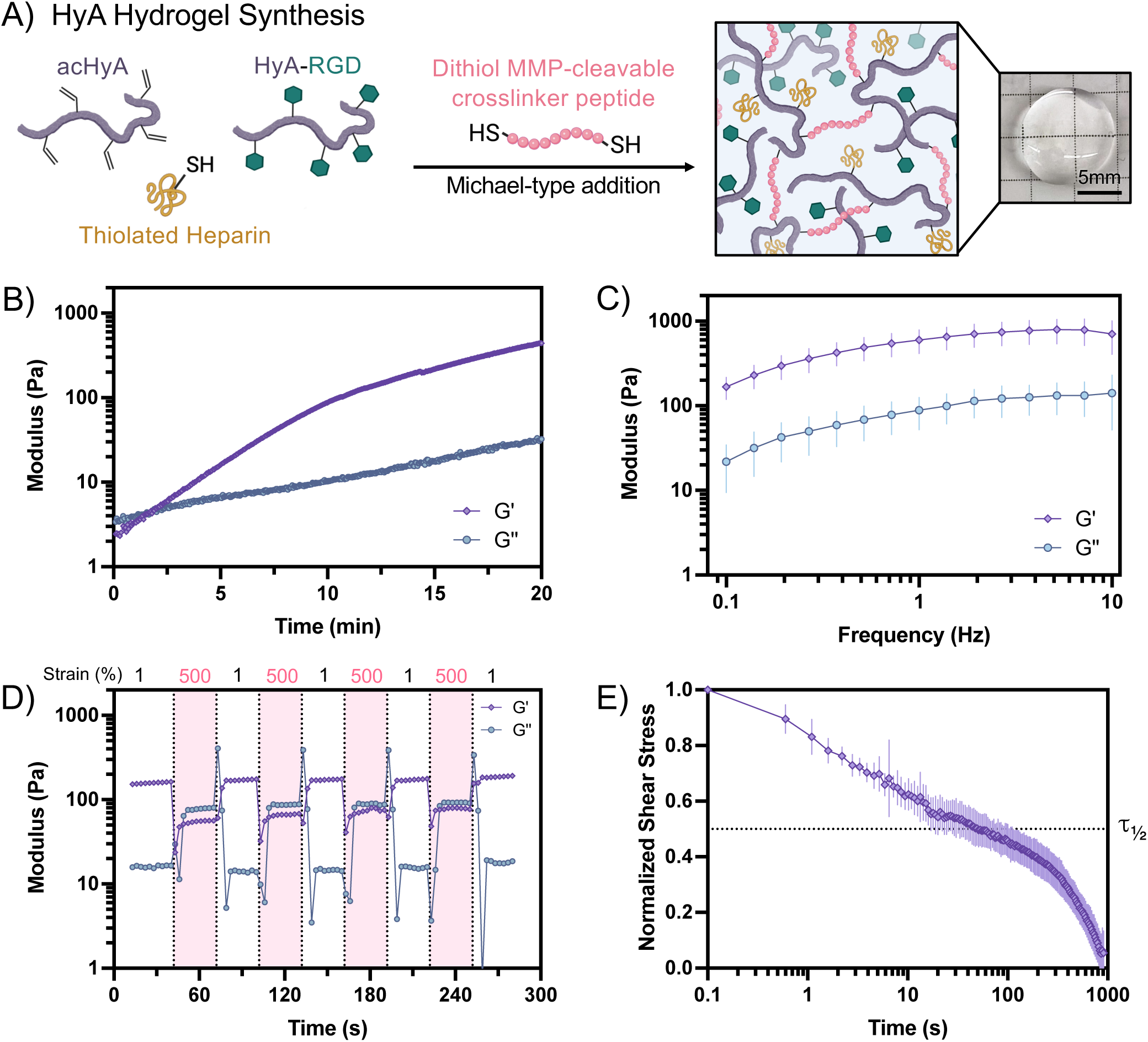
HyA Hydrogel Fabrication and Property Assessment. A) Schematic representation of the formation of HyA hydrogels for rotator cuff repair treatment. Combination of precursor molecules with the dithiol crosslinker peptide initiates gelation via Michael-type addition to form a network. B) Representative gelation (time) sweep of storage (G’, purple diamond) and loss (G’’, blue circle) moduli of the hydrogel immediately after mixing the macromer precursor and crosslinker solutions. C) Storage (G’, purple diamond) and loss (G’’, blue circle) moduli of fully crosslinked HyA hydrogels over a range of frequency values from 0.1 to 10 Hz at 1% strain. Values represent mean ± standard deviation (n=7). D) Representative dataset of storage (G’, purple diamond) and loss (G’’, blue circle) moduli values of a fully crosslinked HyA hydrogel at alternating high-low strain intervals (500% and 1%, respectively) at a frequency of 1 Hz. E) Normalized shear stress of HyA hydrogels after an applied constant 15% strain. Values represent mean ± standard deviation (n=4).

The purpose of this study is to validate the regenerative potential of this HyA hydrogel in a murine model of rotator cuff injury and delayed repair. We assessed material properties of the HyA hydrogel that affects its behavior during implantation/injection and within the regenerative niche including material stiffness, frequency-dependent behavior, self-healing, swelling dynamics, and resulting mesh size. Furthermore, the ability for this hydrogel to sustain the survival, proliferation, and differentiation status of hFAPs was assessed over time in 3D culture. Cell-hydrogel constructs were assessed for expression of a BAT-specific marker, Uncoupling Protein 1 (UCP1), as well as a marker specific to myofibroblasts, α-smooth muscle actin (αSMA). Finally, an *in vivo* study using a delayed rotator cuff repair murine model was conducted to assess post-tear recovery using the HyA hydrogel versus a PBS control. Because fibrosis and FI are hallmarks of muscle degeneration, histology was completed 6-weeks post repair to measure the fat and fibrosis indices of each treatment group. Immunocytochemistry of histological sections revealed a significant degree of vascularization as measured by staining for CD31+ endothelial cells, in alignment with results from prior studies that suggest highly angiogenic properties of our HyA hydrogel.^[25–30]^ We hypothesize that this HyA hydrogel supports promyogenic phenotypes for encapsulated hFAPs. Further, we hypothesize that the hydrogel will reduce FI and fibrosis *in vivo* while promoting vascularization of the affected region.

## 2. Results

### 2.1. HyA Hydrogel Formation and Rheological Properties

Oscillatory rheology was performed to assess the viscoelastic properties of the HyA hydrogel, including gelation kinetics, stress relaxation, and frequency- and strain-dependent behavior of the network. The crossover point of the storage (G’) and loss (G’’) moduli was ∼ 2 minutes, marking the onset of gelation (**Figure 2B**). By 20 minutes, G’ plateaued to ∼ 400 Pa. An increase in both G’ and G’’ values were observed with increased frequency of oscillation, with an initial G’ of approximately 180 Pa at 0.1 Hz that increased steadily up to ∼ 800 Pa at 5 Hz, after which both G’ and G’’ experienced a drop down to 700 Pa by 10 Hz (**Figure 2C**). The observed frequency-dependent changes in hydrogel stiffness inspired investigation into potential strain-dependent properties of the system. At low strain (1%), the hydrogel exhibited a G’ of approximately 200 Pa and G’’ of 15 Pa; these stiffness values changed drastically upon the transition to high strain (500%), as G’ decreased to approximately 60 Pa and G’’ increased to ∼ 100 Pa with a notable transition to G’’ > G’ (**Figure 2D**). The self-healing properties of the HyA hydrogel were demonstrated by its return to initial modulus values upon reversal to low strain. This behavior was repeatable for up to four cycles of alternating high and low strains. Finally, the HyA hydrogel exhibited stress relaxation with a *τ_½_* value of ∼ 60 s, relaxing to a stress as low as 0.1 times its initial value by ∼ 600 s (**Figure 2E**).

### 2.2. HyA Hydrogel Swelling Properties Remain Consistent Across Varying Solvents

The different media used to swell hydrogel samples increased the hydrogel mass but were not statistically significant between conditions (**Figure 3A**). Hydrogels swollen in PBS exhibited a *Q_m_* value of 34.7 ± 4.6, whereas those swollen in F-10 and F-10 + FBS were 39.9 ± 1.7 and 43.4 ± 3.7, respectively (**Figure 3B**). Statistically, there was no significant difference in *Q_m_* between the three media. *Q_v_* values equated to 42.5 ± 5.6, 48.8 ± 2.1, and 53.1 ± 4.6 for PBS, F-10, and F-10 + FBS swollen gels, respectively (**Figure 3C**). There was no statistically significant difference between *Q_v_* values, indicating no significant difference in mass or volumetric swelling for HyA hydrogels in the different media.

**Figure 3:**
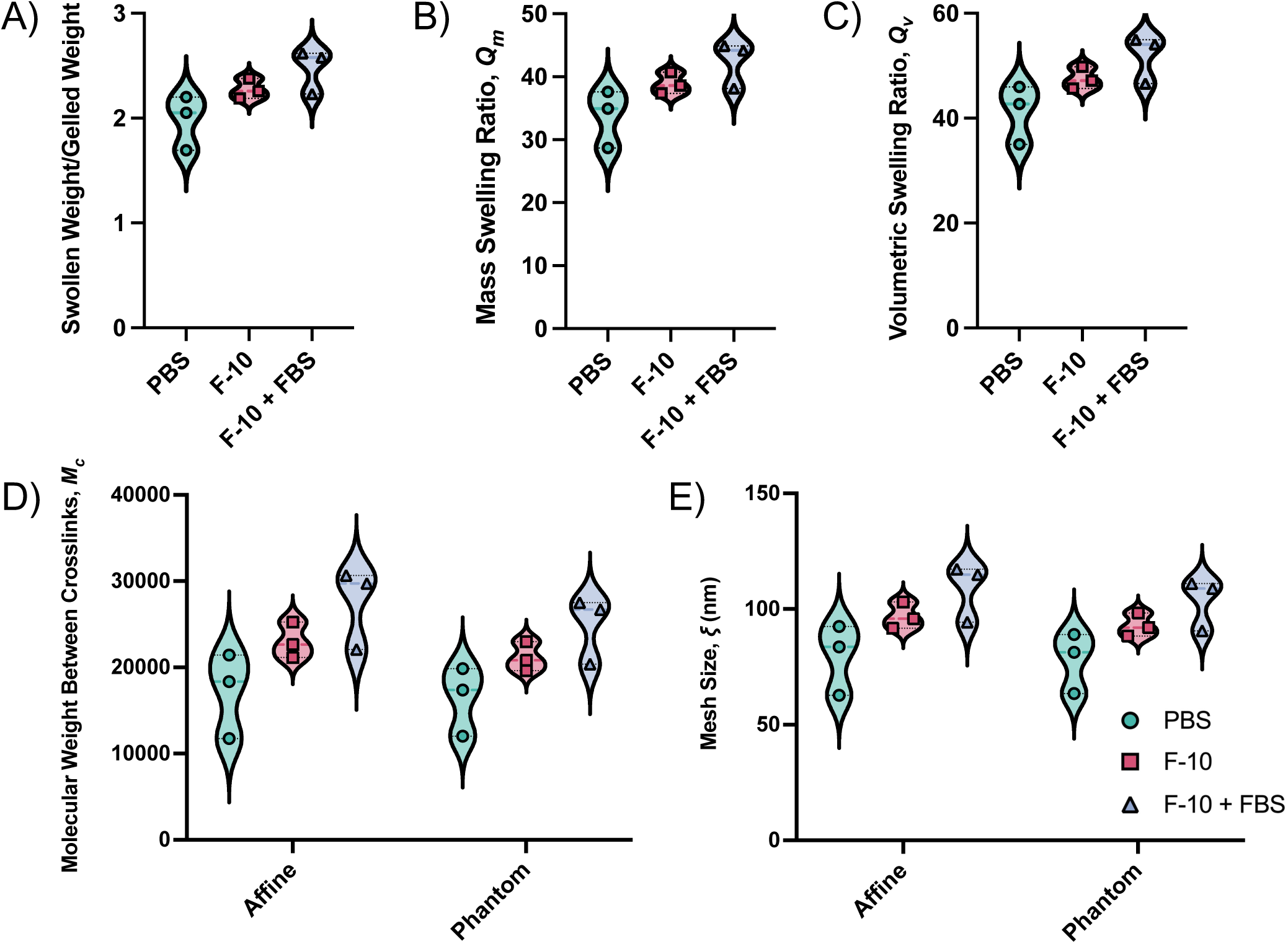
Swelling of HyA Hydrogels in Different Solvents. A) Calculated ratio of swollen to gelled (pre-swelled) hydrogel weight in each solvent tested: PBS, F-10 base media, and F-10 + 10% FBS. B) Calculated mass swelling ratios for hydrogels swollen in each solvent. C) Calculated volumetric swelling ratios for hydrogels swollen in each solvent. D) Approximate molecular weight between crosslinks (Mc) calculated via the affine and phantom network models for hydrogels swollen in each solvent. E) Approximate mesh size (ξ) calculated via the affine and phantom network models for hydrogels swollen in each solvent. Statistical analysis of all datasets was completed using a one-way ANOVA with Tukey’s multiple comparisons test (n=3).

*M_c_* for each swelled hydrogel system was calculated via the theoretical affine (**Eq. 2**) and phantom (**Eq. 3**) network models (**Figure 3D**) to consider two opposite extremes of network deformation theory. *ξ* was finally calculated using *M_c_* from each model in **Eq. 4**. According to the affine model, resulting mesh sizes for hydrogels swelled in PBS, F-10, and F-10 + FBS were 79.6 ± 15.3 nm, 96.9 ± 5.8 nm, and 108.8 ± 12.6 nm respectively. Largely in agreement the phantom model the average mesh sizes equated to 77.9 ± 13.1 nm, 92.9 ± 5.1 nm, and 103.5 ± 11.2 nm, respectively (**Figure 3E**). There was no significant difference in resulting *ξ* between the swelling conditions tested for the HyA hydrogel.

### 2.3. HyA Hydrogels Promote hFAP Survival, Proliferation, and BAT Differentiation in 3D Culture

HyA hydrogels were assessed for their ability to support hFAP survival, proliferation, and bFAP differentiation. hFAP viability was high 24 hours post-encapsulation and remained high 2 weeks into 3D culture experiments (**Figure 4A**). hFAP proliferation steadily increased over time, reaching an approximately 3-fold increase in cell number by 1 week in culture and nearly 5-fold increase by 10 days (**Figure 4B**). hFAP morphology and differentiation trajectory were visualized and quantified after 1, 4, 7, and 10 days of culture (**Figure 4C**). hFAP spreading within the matrix was evident with a significant increase in the average cell volume over time (D1 vs. D10, *p* < 0.0001, **Figure 4D**). UCP1 expression increased steadily over the 10-day period in these hydrogels (D1 vs. D10, *p* < 0.0016, **Figure 4E**), while αSMA expression significantly decreased (D1 vs. D10, *p* < 0.0001, **Figure 4F**).

**Figure 4:**
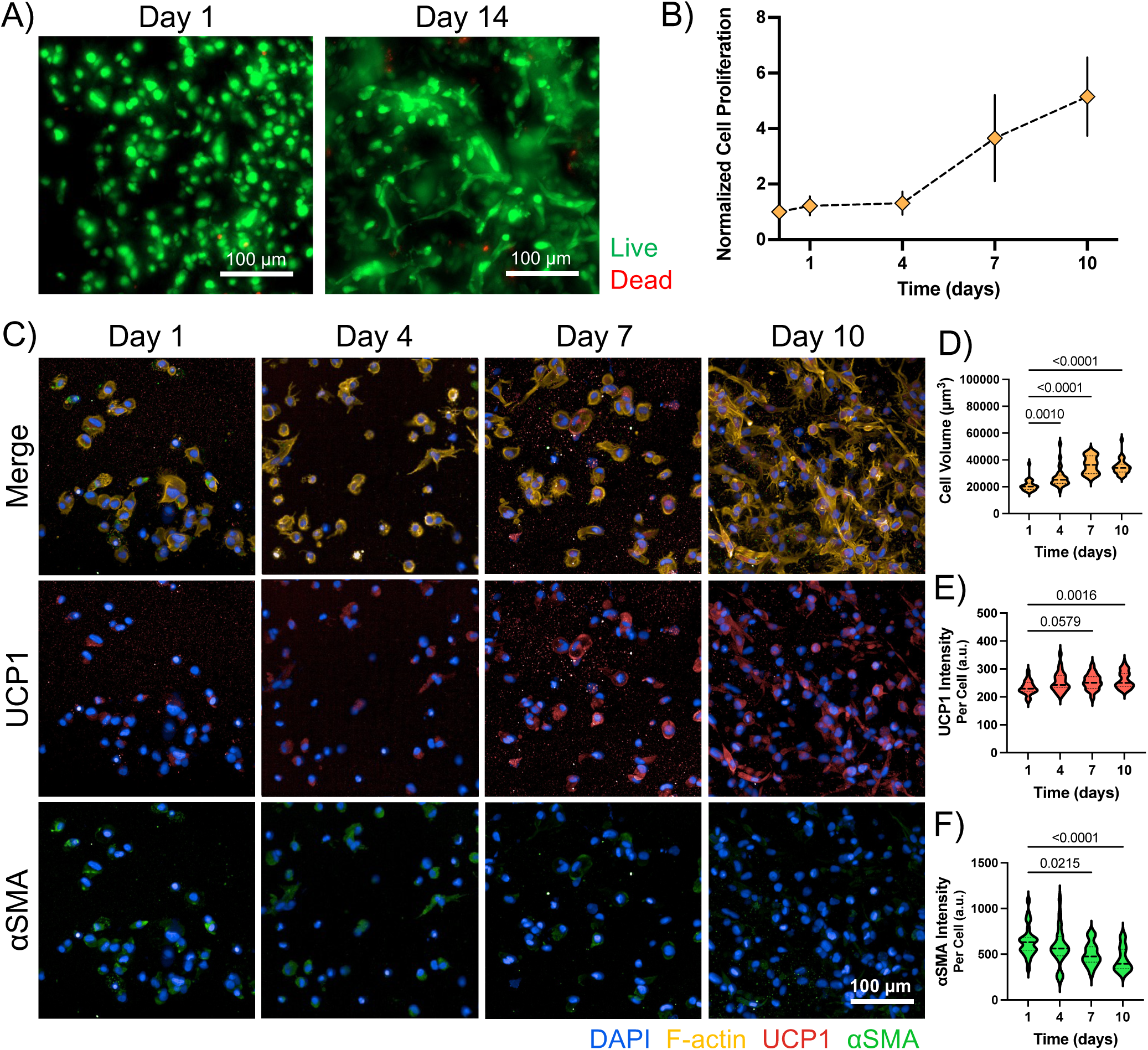
hFAP Survival, Proliferation, and Differentiation in HyA Hydrogels over Time. A) Representative images of HyA-encapsulated hFAPs stained for live (green) and dead (red) markers after 1 day and 14 days in 3D culture. B) Relative cell proliferation (normalized to day 0) as measured by Alamar Blue assay (n=4) over the 10-day period. C) Representative confocal images of hFAP cells in 3D HyA hydrogel culture over 2 weeks. D) Quantification of cell volume at 1-, 4-, 7-, and 10-days post encapsulation. E) Quantification of UCP1 (BAT marker) protein intensity at 1-, 4-, 7-, and 10-days post encapsulation. F) Quantification of αSMA (fibrotic marker) protein expression at 1-, 4-, 7-, and 10-days post encapsulation. Statistical analysis of immunostaining quantifications was completed using a one-way ANOVA with Dunnett’s multiple comparisons test (n=25).

### 2.4. Protein Expression and Cytokine Secretion by hFAPs in HyA hydrogels

The pro-angiogenic outcomes historically achieved using this hydrogel system with a variety of cell types inspired investigation into hFAP angiogenic behavior while encapsulated in the HyA hydrogel.^[25–29]^ hFAPs were isolated from hydrogels at 1, 8, and 14 days post-encapsulation and assessed for the expression of 55 angiogenesis-related factors (**Figure 5A**). Among the 55 proteins tested, 21 experienced at least a 5-fold increase from day 1 to day 14, with another 18 proteins at least doubling in expression over the 2-week period (**Figure 5B**). Among the most highly upregulated proteins was Activin A with a 265-fold increase in expression, followed by FGF-7 increasing by 114-fold (**Figure 5C**). Prolactin, angiostatin/plasminogen, and MCP-1 were considerably increases in this time, between 22 and 35 times their initial expression levels on D1, while 10 additional proteins including FGF-basic and leptin increased by 6- to 12-fold. By day 14, hFAPs exhibited the overall highest expression of activin A, followed by DPIV, Serpins F1 and E1, FGF-basic, and urokinase plasminogen activator (uPA) (**Figure 5D**). As an additional assessment of regenerative hFAP behavior within the gels, the level of anti-inflammatory cytokine IL-10 in the sample media was assessed at 1- and 7-days post-encapsulation. A significant increase in IL-10 was observed over the 1-week period, approximately 6 times its initial expression from 5 to 30 ng/mL (*p* = 0.0041) (**Figure 5E**).

**Figure 5:**
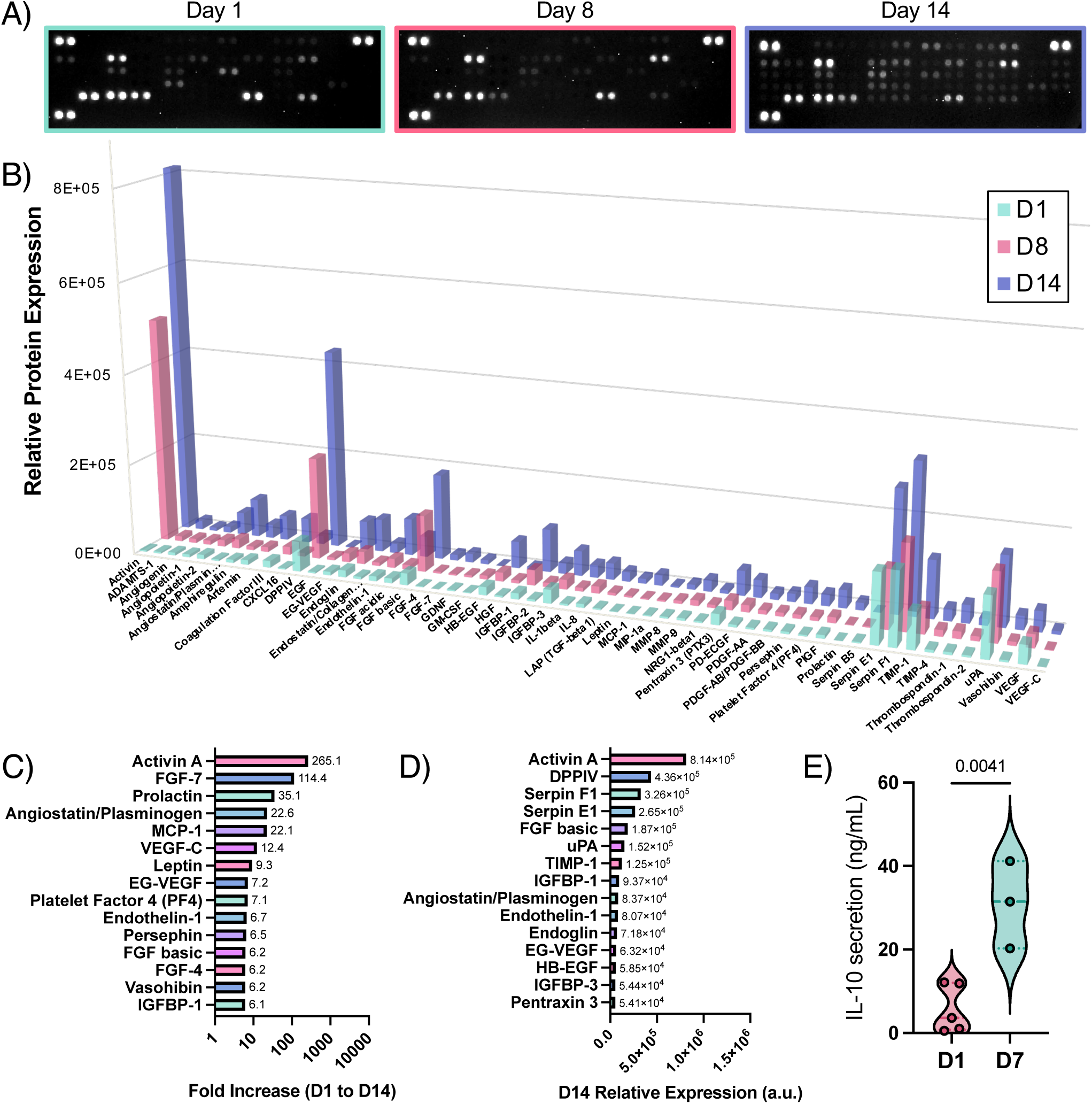
Protein Expression & Cytokine Secretion by hFAPs in HyA Hydrogels. A) Chemiluminescent arrays were used for protein samples collected from hFAPs 1-, 8-, and 14-days post-encapsulation to assess the relative expression of 55 different angiogenic proteins. B) Quantified relative protein expression of angiogenesis-related proteins at each time point. C) The top 15 proteins as ranked by fold-increase from day 1 to day 14 after encapsulation. D) The top 15 proteins as ranked by the highest overall expression level at day 14 after encapsulation. E) IL-10 secretion level in the media surrounding cell-gel constructs at 1- and 7-days post-encapsulation. Day 1 (n=5) and 7 (n=3) time points were statistically compared using a Student’s t-test.

### 2.5. Hydrogel Implantation Reduces Muscle Atrophy

Assessment of the HyA hydrogel *in vivo* was completed using a rigorous delayed repair murine model of rotator cuff injury. 6-weeks after creation of the injury, animals received surgical repair in addition to the HyA hydrogel or PBS control treatment (**Figure 6A**). The wet muscle weight of right-sided supraspinatus (RSS) from injured/repaired limbs when compared to that of the contralateral, non-injured left-sided supraspinatus (LSS) control showed that hydrogel treatment led to a relative increased retention in muscle weight when compared to PBS treatment 6 weeks post-injury (-67.66% in PBS vs -34.66% in gel, normalized to LSS, *p* = 0.0032) (**Figure 6B**). Laminin staining (**Figure 6E**) revealed a decrease in the average myofiber cross-sectional area (CSA) after injury in PBS treated limbs compared to contralateral controls (2206 μm^2^ in PBS LSS vs 1510 μm^2^ in PBS RSS, *p* = 0.0006), while hydrogel treated CSA levels were comparable to that of the alternate control limb (2148 μm^2^ in gel LSS vs 2096 μm^2^ in gel RSS, *p* = 0.9896) (**Figure 6C**). The average CSA resulting from PBS treatment is also significantly lower than that of hydrogel treated limbs (1510 μm^2^ in PBS RSS vs. 2096 μm^2^ in gel RSS, *p* = 0.0046). When normalized to non-injured LSS, hydrogel treatment led to a relative decrease in CSA reduction compared to PBS treatment 6 weeks post-injury (-31.33% in PBS vs -1.24% in gel, normalized to LSS, *p* = 0.0005) (**Figure 6D**).

**Figure 6:**
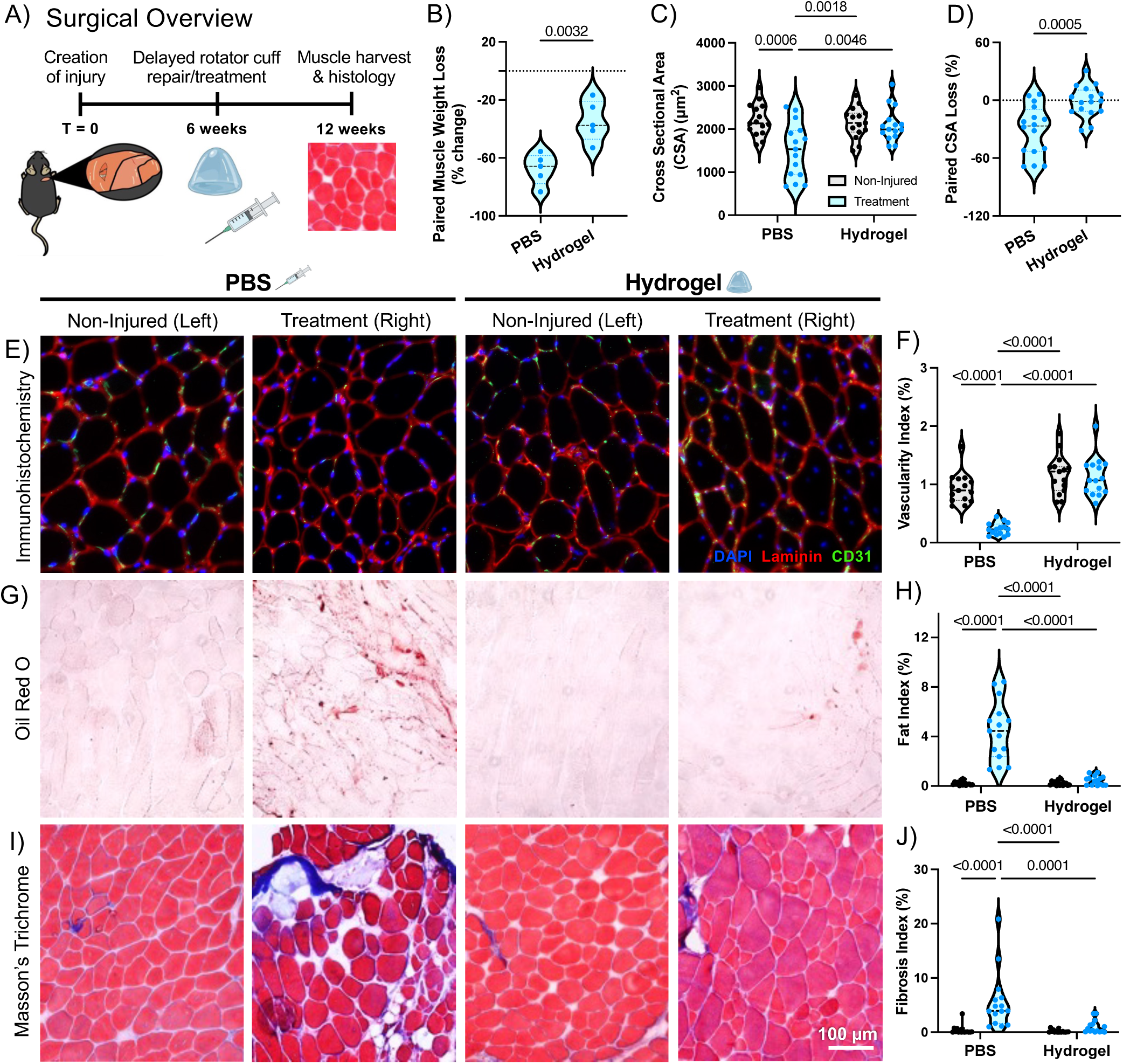
*in vivo* Assessment of Delayed Rotator Cuff Repair Using HyA Hydrogel. A) Schematic of experimental timeline. Rotator cuff injury was induced by surgical tenotomy and denervation. After 6 weeks, the repair surgery was performed with either hydrogel or PBS injection into the muscle belly. 6 weeks post-repair, muscle from the treated limb (right) and non-injured contralateral control limb (left) was harvested and assessed for quality, weight, and sectioned for histological staining. B) Paired muscle weight loss plotted as a percent change from control, non-injured animal muscle. PBS and hydrogel treatments (n=5) were compared using a Student’s t-test. C) Quantification of cross-sectional area (CSA) as measured by calculation of area of bundles contained within laminin-stained regions (red) of stained tissue sections. D) Paired CSA loss plotted as a percent change from control, non-injured animal muscle. PBS and hydrogel treatments (n=15) were compared using a Student’s t-test. E) Immunocytochemistry representative images of sections from each treatment group to assess for vascularization and muscle fiber structure/area. From left to right: non-injured (PBS treatment contralateral control), injury with PBS treatment, non-injured (hydrogel treatment contralateral control), injury treated with HyA hydrogel. F) Quantification of vascularization as measured by calculation of % area of CD31+ stained regions (green) of stained tissue sections. G) Oil Red O staining representative images of histological sections from each treatment group to assess for fatty infiltration. H) Quantification of fat index as measured by calculation of % area of Oil Red O positive stained regions (red) from histology. I) Masson’s Trichrome staining representative images of histological sections from each treatment group to assess for fibrosis. Scale bar = 100 µm. J) Quantification of fibrosis as measured by calculation of % area of collagen positive stained regions (purple) from histology. CSA and immunostaining quantifications were statistically analyzed using a two-way ANOVA with Tukey’s multiple comparisons test (n=15).

### 2.6. Hydrogel Implantation Preserves Muscle Vascularity

Immunohistochemistry staining of CD31 (**Figure 6E**), representative of surface of endothelial cells and vascularization, showed a marked decrease in vascularity after injury in PBS treated limbs compared to contralateral LSS (0.932% in PBS LSS vs 0.235% in PBS RSS, *p* < 0.0001), but no significant difference between hydrogel treated RSS and the control limb (1.168% in gel LSS vs 1.129% in gel RSS, *p* = 0.9793) (**Figure 6F**). Direct comparison between PBS treated and hydrogel treated limbs indicated a significant increase in vascularity in the hydrogel treated group compared to PBS treatment (0.235% in PBS RSS vs. 1.129% in gel RSS, *p* < 0.0001).

### 2.7. Hydrogel Implantation Significantly Reduces Fibrosis and Fatty Infiltration

As indicated via Oil Red O staining (**Figure 6G**), PBS treated mice experienced a significant increase in fat index in repaired RSS muscle compared to control LSS (0.204% in PBS LSS vs 4.442% in PBS RSS, *p* < 0.0001), while comparable FI was observed between hydrogel RSS and hydrogel LSS; 0.221% in gel LSS vs 0.452% in gel RSS, *p* = 0.9535) (**Figure 6H**). Comparison of RSS injured limbs between treatments showed decreased fat index in the hydrogel treated group compared to PBS (4.442% in PBS vs 0.452% in gel, *p* < 0.0001).

Masson’s trichrome staining (**Figure 6I**) indicated that for PBS treated mice, there was a significant increase in fibrotic area in injured RSS compared to control LSS, while comparable fibrosis was observed between hydrogel RSS and hydrogel LSS (0.351% in PBS LSS vs 5.618% in PBS RSS, *p* < 0.0001; 0.131% in gel LSS vs 0.967% in gel RSS, *p* = 0.9569) (**Figure 6J**). Comparison of right-sided injury showed decreased fibrotic index in right-sided hydrogel group compared to PBS (5.618% in PBS RSS vs 0.967% in gel RSS, *p* = 0.0001).

## 3. Discussion

Rotator cuff tears are one of the most common musculotendinous injuries encountered by physicians, yet they have remarkably poor post-injury outcomes. Patient outcomes could be significantly augmented with promyogenic therapeutics that properly modulate the regenerative response. Our HyA hydrogel formulation is well-suited for this application, having demonstrated the ability to augment CD31 expression of vascular cells *in vitro*^[22]^ ^Jha, Tharp, 2015; [23]^ ^Jha, Tharp, 2016; [24]^ ^Jha, Mathur, 2015; [25]^ ^Browne, Jha, 2018; [26]^ ^Browne, Hossainy, 2020^ and reduce muscle weight loss while increasing functional recovery and vascularization after VML injury *in vivo*.^[30]^ In this study, we aimed to assess the capacity for this hydrogel to be used as an adjunct to rotator cuff surgical repair to prevent the fibrosis and FI typically observed after chronic injury.

Biomaterial mechanical properties are crucial to determining the success of a therapeutic, from ease of clinical administration to the biological response invoked by the implant. Hydrogel properties like gelation time, stiffness, and swelling dynamics are crucial to consider when designing a therapeutic such as *in situ* hydrogels. The solution precursors are relatively low viscosity, making them well-suited for injectable systems and easier to work with than many high-viscosity alternatives. As indicated by the sol-gel transition, the system forms a gel after approximately 2 minutes of crosslinking but continues to stiffen for another 20 mins (**Figure 2B**), providing ample working time for clinical delivery. Nonetheless, gelation time is rapid enough to ensure the mechanical integrity of the hydrogel at time of implantation, preventing material drift from the site of injury.

Viscoelasticity is an established property of native tissues shown to regulate critical cellular and regenerative processes,^[31,32]^ making it a key parameter of investigation in this study. In particular, FAP activation and differentiation have been previously shown to be dependent on matrix mechanical properties like stiffness,^[33]^ reinforcing the need for integration of appropriate viscoelastic cues into materials designed to influence FAP trajectory. Frequency-dependent behavior is one such property associated with viscoelastic materials, with higher frequencies generally correlating with more elastic behavior like increased G’. This behavior is observed in the HyA hydrogel system, with an initial G’ of approximately 180 Pa that steadily increases with frequency up to 4 Hz (**Figure 2C**). However, at this point, both G’ and G’’ begin to drop up to the highest tested frequency. While not expected, this drop may be explained by chain rearrangements and release of entanglements within the polymer network that reduce the system’s ability to store and dissipate energy. At high frequencies, should network deformation occur faster than the polymer chains’ ability to recover from it, the material’s viscous and elastic components may be reduced, as reflected in these lower G’ and G’’ values.

This dynamic frequency-dependent behavior inspired investigation into additional viscoelastic properties like strain dependency. Subjecting the hydrogels to alternating high-low shear strain rates revealed striking phase-change and recovery behavior (**Figure 2D**). Upon transition from 1% to 500% strain, the sol-gel transition indicated by G’’ exceeding G’ suggests liquification of the hydrogel, with almost immediate recovery to G’ > G’’ after reversal back to 1% strain. Impressively, the hydrogel sustains at least 4 high-low strain cycles with a return to approximately its initial G’ and G’’ values, indicating repeatable cycles of solid-like to liquid-like form at shear strain extremes with minimal permanent deformation. This behavior may be explained by system-wide macromer chain rearrangements. Given the relatively high molecular weight of the system that consists of an average of 1,319 monomers per chain, physical (and reversible) crosslinks like entanglements, hydrogen bonding, and other chain interactions are prevalent and contribute to the overall elastic component of the material. Furthermore, HyA-RGD chains—comprising 60% of the system’s total macromer—experience almost complete substitution of the system’s acrylate groups as demonstrated via NMR spectra and lack of ability to form a gel in the presence of excess crosslinker. Thus, there are few acrylate groups remaining available for chemical conjugation into the overall network. This semi-interpenetrating network (IPN)-like structure enhances the ability for chains to reorganize under stress or deformation. Thus, despite large overall strains or oscillation rates, the non-permanent and reversible nature of these interactions allow for the transition to and from the gel- and fluid-like states in addition to enabling self-healing behavior. Also consistent with this interpretation is the rapid and near-complete relaxation of stress after a constant applied 15% strain to the hydrogel (**Figure 2E**). The hydrogel exhibits a *τ_½_* value of approximately 6 minutes, comparable to the stress relaxation times of many native human tissues and significantly faster than that of many covalently crosslinked hydrogels.^[34]^ In addition to the relatively low *τ_½_*, the system’s ability to reach nearly 10% of its initial stress value could also be explained by polymer chain sliding and disruption of physical crosslinks to dissipate the applied stress.

As hydrogels are predominantly comprised of a liquid component, their swelling behavior—and therefore material properties—are impacted by the effective solvent within the surrounding environment. Therefore, the effects of PBS, F-10 base cell culture media, and F-10 + 10% FBS (standard hFAP culture media) as swelling agents were investigated. While PBS or similar, serum-free solvents are typically used in material property characterization, assessment of our HyA hydrogel in F-10 media, particularly that supplemented with FBS, provided a more physiologically relevant (i.e. implanted hydrogel in contact with blood and protein-rich bodily fluids) measurement of critical material parameters. Recording the mass of hydrogels before and after complete swelling in one of the three solvents enabled calculation of key swelling parameters, *Q_m_* and *Q_v_*, for each sample group (**Figure 3A-C**). Somewhat surprisingly, there was no significant difference in either metric between hydrogels swollen in each solvent. While the diffusion-based influx of proteins and other molecules in cell culture media and serum would be expected to increase hydrogel swelling compared to the PBS-only condition, the observed behavior may be accounted for by the presence of entanglements between HyA polymer chains that resist swelling-induced network deformation and expansion. This theory is reflected in the approximated values for hydrogel mesh size determined via two different network models used for material properties prediction. Specifically, the affine (**Equation 2**) and phantom (**Equation 3**) network models enable the calculation of *M_c_* using swelling data collected in this study in addition to known material and solvent parameters (**Figure 3D**). Via the affine model, the default polymer network model, crosslinks are assumed to be rigidly fixed to the gel body and move in proportion to macroscopic deformation, entirely accounted for by chain uncoiling and stretching. Using affine-predicted *M_c_*, *ξ* was calculated as in **Equation 4** for hydrogels swollen in each solvent (**Figure 3E**). No significant difference in mesh size was observed between hydrogels in each condition, which range from 80-110 nm in size. A similar trend was observed using the phantom network model, which produced mesh size values from 78-105 nm in size for the differently swollen gels. In this model, crosslinks are not fixed but fluctuate around an average position, thereby suppressing chain deformation and reducing the expected mesh size. As the two models are considered opposite extremes on the mesh size calculation spectrum, the use of both approaches provides a relative range in which we can expect our system’s mesh size to fall. Physiologically, this scaffold’s mesh size—and ultimately the porosity within the network experienced by infiltrating biologics—provides ample room for diffusion of nutrients and waste into and out of the system, as well as cell infiltration and ECM deposition by FAPs that rapidly expand and enter a matrix-producing, pro-inflammatory phenotype after injury.^[9]^

Given the multipotent nature of hFAPs and their dynamic orchestration of the regeneration process, assessing their response to the hydrogel *in vitro* provided insight into their possible *in vivo* therapeutic response to implantation. The hydrogel proved to effectively support hFAP survival, growth, and proliferation in 3D culture (**Figure 4A-D**). Cell spreading occurred uniformly throughout the cell-gel constructs. Furthermore, the trends in UCP1 expression and αSMA expression over time are in alignment with what would be expected for promyogenic FAP activation, as BAT expression picks up with a reduction of ECM-producing fibroblasts (**Figure 4C-F**). While fibrotic expression was not anticipated at baseline, these hFAPs are patient-derived and often have some degree of heterogeneity,^[35]^ which means some encapsulated FAPs may have been trending in a fibrotic lineage at the time of encapsulation. The ability for the HyA hydrogel to reduce fibrotic behavior is indicative of its promyogenic capacity and aligns well with the aim to minimize fibrosis and FI in this injury model. It has been reported using collagen hydrogels that increased substrate stiffness up to 50 kPa correlates with increased expression of αSMA in FAPs, even more so when cultured on rigid plastic.^[33]^ Perhaps the change from rigid TCPS to the softer hydrogel environment upon encapsulation may have contributed to the reduction of this phenotype.

Investigation into hFAP protein expression over time was also conducted to elucidate changes over the course of the encapsulation. Historically, excellent vascularization has been achieved *in vitro* and *in vivo* using this HyA hydrogel.^[25–29]^ hFAPs were therefore isolated from cell-gel constructs at different time points to conduct an angiogenesis-specific proteome analysis. As measured both by fold increase in expression over the 2-week period and overall expression level at day 14, Activin A was ranked number one for both categories (**Figure 5C, D**). Activin A has been shown to stimulate endothelial cell proliferation, migration, and tube formation, directly supporting angiogenesis and tissue vascularization.^[36]^ While Activin A has also been associated with increased expression of ECM proteins in FAPs that is typically associated with fibrosis,^[11]^ production of ECM within an acceptable range is a necessary component of regeneration and will occur as the hydrogel is degraded by cell-secreted MMPs. A variety of other pro-angiogenic proteins were shown to be upregulated over time including FGF basic,^[37]^, FGF-4,^[38]^ VEGF-C,^[39]^ and urokinase-type Plasminogen Activator (uPA).^[40]^

The promyogenic capabilities of this HyA hydrogel were further highlighted in a rigorous delayed rotator cuff repair model designed to mimic a clinically relevant scenario where fibrosis, FI, and atrophy have already developed. Following the development of a chronic tear 6 weeks post-injury, the tendon was repaired and muscle treated with HyA hydrogel or PBS. FI and fibrosis are hallmark indicators of irreversible muscle pathogenesis, impairing muscle function, negatively affecting muscle regeneration after injury, and increasing muscle susceptibility to re-injury. Chronic rotator cuff tears also induce muscular atrophy, characterized by a decrease in myofiber size and overall muscle volume loss. Rotator cuff tears have been shown to induce atrophy-related genes, increasing the severity of muscle degeneration.^[41]^ Treatment with the HyA hydrogel minimized fibrosis and fat indices nearly indistinguishable from those of non-injured contralateral control limbs. The near absence of these anti-myogenic features is remarkable and indicative of a potential biological landscape change fostered by the introduction of the HyA hydrogel. Furthermore, we found that hydrogel post operatively reduces the degree of atrophy and increases muscle cross-sectional area in this model, both important markers of muscle size. We believe one of the mechanisms that aids in our hydrogel to achieve such outcomes is its ability to induce FAPs into its promyogenic beige fat phenotype.^[42,43]^ Prior studies have shown when beige FAPs are administrated to a chronic rotator cuff injury, it leads to decreased FI and fibrosis.^[15,44]^ Our *in vitro* experiments indicated that encapsulated FAPs differentiated into its promyogenic phenotype suggesting this is perhaps the mechanism of action of our hydrogel.

Skeletal muscle is inherently a highly vascular tissue due to the high metabolic demands of contraction, and angiogenic and myogenic processes have been shown to be exhibit cellular crosstalk.^[45]^ Thus, increasing vascular supply to muscle injury may be an approach to improve muscle regeneration.^[46]^ Vessels provide metabolic components and remove biological waste in the muscle environment while serving as highways for various cell types throughout the body, making them indispensable for successful muscle regeneration. Quantification of immunohistochemical (CD31) staining revealed a vascular index for HyA-treated injuries that is significantly greater than the repaired, PBS-treated control and equal to that of contralateral controls, indicating successful vascularization of the regenerated tissue. This correlates well with previous studies in showing the capability of HyA in improving/preserving vascularity.^[30,47]^ Previous studies on vascularity in rotator cuff has shown less vascular microcirculation in degenerative conditions.^[48,49]^ While there is little information on the relative importance of vascularity in muscle recovery in rotator cuff tears, it stands to reason that improved vascularity is an important aspect of chronic muscle recovery in this state as well.

Collectively our results suggest a hypothesis where the HyA hydrogel leads to successful repair of a chronic rotator cuff injury model by inhibiting FAP differentiation to anti-myogenic phenotypes including fibroblasts and white fat, leading to reduced scar tissue and FI buildup, while actively promoting the development of FAPs into promyogenic lineages including bFAPs (**Figure 6**). We believe that our HyA hydrogel helps reshape the landscape of the injured rotator cuff niche by improving overall muscle quality and impacting FAP cell fate to reduce the production of non-contractile tissue. Furthermore, at the time of surgery, the muscle belly of the rotator cuff (supraspinatus and infraspinatus) is accessible through arthroscopic portals, suggesting that targeted delivery of injectable scaffolds to improve muscle quality could be achievable with minimal patient morbidity. Taken together, these results may form the foundation of future rotator cuff repair therapeutics while serving as a potential testbed for further FAP investigation.

**Figure 7:**
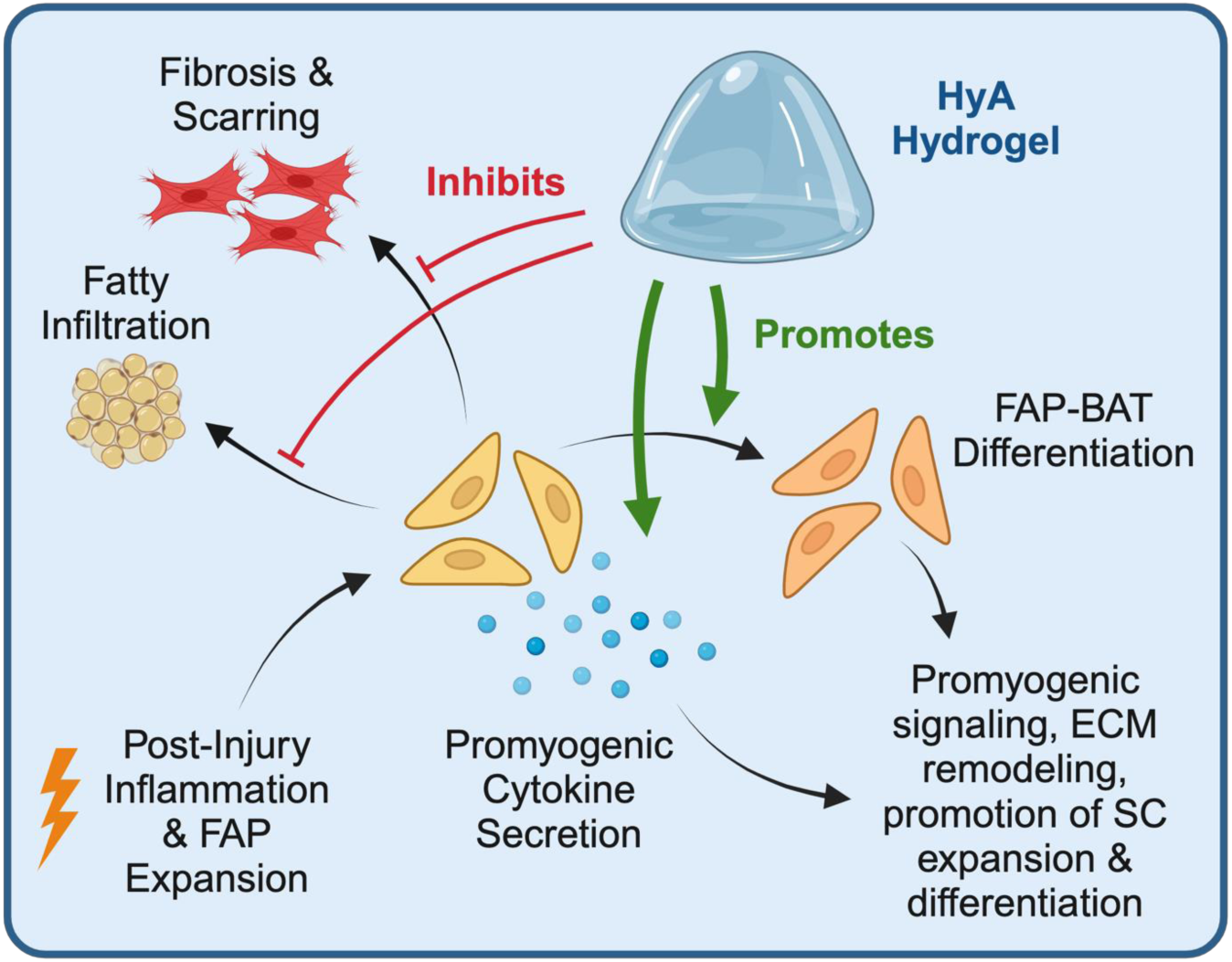
Proposed Mechanism of HyA Hydrogel. After injury, FAP expansion is promoted by inflammation and other signals. FAPs may then differentiate into myofibroblasts (fibrosis), white fat (FI), or pro-myogenic brown fat that encourages downstream regeneration. The results from this study suggest that the HyA hydrogel has an inhibitory effect on the production of scar tissue and FI after injury, while promoting pro-myogenic bFAP differentiation.

## 4. Conclusion

Our HyA hydrogels exhibit viscoelastic properties well-suited for muscle tissue regeneration applications, such as rotator cuff repair. Our *in vitro* work demonstrated the ability of this HyA hydrogel to support FAP proliferation, spreading, and BAT differentiation while increasing promyogenic cytokine secretion over time. *In vivo*, application of our HyA hydrogel to a delayed repair murine rotator cuff model reduced fibrosis, FI, and muscle atrophy while supporting vascularization of injured muscle tissue, ultimately leading to regeneration of muscle tissue. Together, our results suggest that hydrogel-mediated FAP activity may contribute to the positive regenerative outcomes observed after treatment. The materials developed during these studies provide the foundation for both *in situ* and *in vivo* biomaterial therapy approaches for treatment of rotator cuff and other musculoskeletal injuries in the clinic. Understanding the biophysical and material parameters that influence bFAP differentiation enable their translation into more effective therapeutics for muscle regeneration and beyond.

## 5. Experimental Section

### Materials

Hyaluronic acid (HyA, Sodium Hyaluronate Powder, 500 kDa) was purchased from Lifecore Biomedical (Chaska, MN). Adipic acid dihydrazide (ADH), 1-Hydroxybenzotriazole hydrate (HOBt), sodium hydroxide (NaOH), and hydrochloric acid (HCl) were purchased from Sigma-Aldrich (Milwaukee, WI). 1-Ethyl-3-[3-(dimethylamino)propyl] carbodiimide (EDC), N-Acryloxysuccinimide 99% (NAS), tris(2-carboxyethyl)phosphine (TCEP), and triethanolamine-buffer (TEOA; 0.2 M, pH 8.0) were purchased from Fisher Scientific (Hampton, NH). Dimethyl sulfoxide (DMSO), N-Acryloxysuccinimide (NAS), and ethanol were obtained from Fisher Scientific (Waltham, MA). Dialysis membranes (10,000 MWCO, SpectraPor Biotech RC) were purchased from Spectrum Laboratories (Rancho Dominguez, CA). Thiolated heparin was purchased from Scientific Protein Laboratories (SPL) (Waunakee, WI). The MMP-degradable cross-linker peptide (CQPQGLAKC) was synthesized by CPC Scientific (Sunnyvale, CA) and bsp-RGD(15) adhesion peptide (Ac-CGGNGEPRGDTYRAY-NH2) was synthesized by United BioSystems (Herndon, VA). All chemicals were used as received. All cell culture reagents were purchased from Invitrogen (Carlsbad, CA). All reactions were completed using MilliQ synthesis-grade ultrapure water (Milli-Q IQ 7000 system).

### Synthesis of Acrylated HyA

HyA-based hydrogels were synthesized using previously reported methods.^[25–30,43]^ Briefly, the HyA derivative carrying hydrazide groups (HyA-ADH) was synthesized by the addition of 30 M excess of ADH to HyA in MilliQ water (200 mL, 3 mg/mL). Solution pH was adjusted to 6.8 using 0.1 M NaOH and 0.1 M HCl. EDC (3 mmol) and HOBt (3 mmol) were dissolved separately in a 1:1 DMSO:water solution and added to the HyA solution sequentially. The pH was maintained at 5.0 for 2 h using 0.1 M NaOH and 0.1 M HCl, after which the solution was neutralized and exhaustively dialyzed against MilliQ water, precipitated in 100% ethanol, and re-dissolved in 200 mL MilliQ water. NAS (1.4 g) was subsequently reacted with the HyA-ADH solution in the absence of light to generate acrylated HyA (acHyA). After 24 hours, the product was exhaustively dialyzed against MilliQ water, sterile filtered through 0.22 µm Millipore Steriflips, and lyophilized until completely dried. Using a previously described analysis,^[50]^ proton (1H) NMR spectra were recorded in D_2_O on a Bruker NEO-500 NMR spectrometer, with an acceptable acrylation range of 20-25% of total monomers (**SI Figure 1**).

### Generation of HyA-bsp-RGD(15)

The HyA-peptide derivative was synthesized by reacting the peptide sequence Ac-CGGNGEPRGDTYRAY-NH2 (bsp-RGD(15), 50 mg) with acHyA solution (125 mg, 50 mL MilliQ water) at 30°C overnight. The peptide was pretreated with excess TCEP for 1.5 hours to reduce any disulfide bonds formed between cysteine thiol groups. The HyA-bsp-RGD(15) product (HyA-RGD) was exhaustively dialyzed against MilliQ water, sterile filtered through 0.22 µm Steriflip vacuum units (Millipore Sigma, #SE1M179M6), and lyophilized until completely dried. 1H NMR spectra were recorded in D_2_O on a Bruker NEO-500 NMR spectrometer to verify successful conjugation of bsp-RGD(15) to the acrylated precursor (**SI Figure 1**). HyA-RGD was validated via ^1^H NMR spectra that demonstrated complete absence of acrylate groups and the presence of tyrosine-associated peaks.

### HyA Hydrogel Formation

Hydrogels were formed as previously described.^[25–30,43]^ Briefly, acHyA (1.33 wt%), HyA-RGD (2.00 wt%), and thiolated heparin (0.03 wt%) were dissolved in TEOA buffer and dissolved at 37°C. Hydrogels were fabricated by mixing a bis-cysteine-terminated, MMP-cleavable crosslinker peptide (CQPQGLAKC) solution (3 mg/50 μL TEOA) with the HyA/HyA-RGD/heparin precursor solution at a 1:6 ratio of crosslinker to precursor by volume.

### Rheology

HyA hydrogel viscoelastic properties were assessed using an oscillatory rheometer (MCR 302, Anton Paar, Ashland, VA). All measurements were taken with the base plate temperature set at 37°C using a humidity-controlled chamber. Hydrogel gelation time was measured using a 25 mm parallel plate measuring system under constant 0.1% strain and 1 Hz frequency for 30 mins. The macromer and crosslinker solutions were mixed and immediately dispensed onto the rheometer for measurement. The crossover point of the storage modulus (G’) and loss modulus (G”) was taken as the gelation time. Viscoelastic properties of hydrogels after complete gelation (1 hour in a 37°C, humidity-controlled incubator) were taken using a sandblasted 8 mm parallel plate measuring system. For frequency sweep tests, samples were subjected to 1% strain and frequency values logarithmically ramping from 0.1 to 10 Hz. Self-healing of the hydrogel was assess via high-low strain measurements. High and low strains of 500% and 1% respectively, were conducted in 30s interval. Stress relaxation measurements were completed by applying an initial constant strain of 15% to the hydrogel system and measuring the shear stress over time. Results are plotted as shear stress values normalized to the initial shear stress value.

### Hydrogel Swelling and Mesh Size Calculations

The swelling properties of HyA hydrogels were determined for different solvents including PBS, base cell culture media (F-10), and media plus 10% fetal bovine serum (FBS). Hydrogels were formed in cylindrical acrylic molds of 8 mm diameter and 1 mm height and gelled for 1 hour at 37°C in a humidity-controlled incubator to ensure complete gelation. The mass of each hydrogel was measured before and after overnight swelling in one of the three solvents. The mass swelling ratio (*Q_m_*) was calculated for hydrogels in each solvent by dividing the mass of the gel in the swollen state by the mass of dry polymer in each sample. *Q_m_* was then used to calculate the volumetric swelling ratio (*Q_v_*) for hydrogels in each solvent using **Equation 1** below, where *ρ_p_* is the polymer (HyA) density and *ρ_s_* is the solvent density:

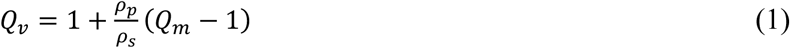

The effect of solvent on the resulting mesh size of the hydrogel was assessed using affine and phantom network models to approximate the effects of swelling. *Q_v_* values for hydrogels swelled in each solvent were first used to calculate the volume fraction of the polymer in the relaxed/pre-swelled state (*v_2,r_*) and the volume fraction of the polymer after swelling (*v_2,s_*). These parameters, along with the number average molecular weight (*M_n_*), dry polymer specific volume 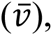 molar volume of solvent (*V_1_*), Flory interaction parameter (*χ_1_*), polymer concentration (*c*), and polymer functionality (*φ*) were used to approximate the average molecular weight between crosslinks 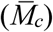 via the affine (**Equation 2**)^[51,52]^ and phantom (**Equation 3**)^[53]^ network models:

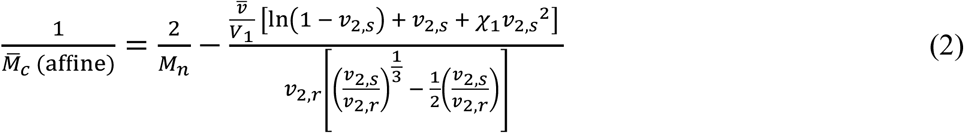

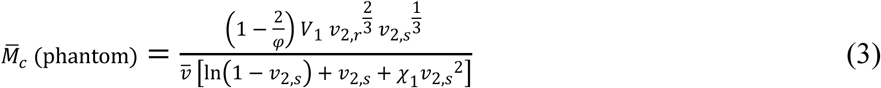

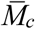 was then used to determine the mesh size (*ξ*) corresponding to each network model using **Equation 4**.^[54]^

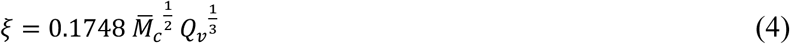

### Isolation of hFAPs from donor muscle tissue

This study was conducted under the approval of the Institutional Review Board (IRB # 18-26000 and 23-40440) at the University of California San Francisco (UCSF). Patient tissue samples were collected with prior informed consent obtained from all participating subjects. All participants were provided with detailed information about the purpose, procedures, risks, and potential benefits of the research before giving their written consent. Human FAPs (hFAPs) were harvested from deltoid muscle biopsies (10 mg samples) performed on patients undergoing arthroscopic rotator cuff repair at UCSF. Muscle was digested with Dulbecco’s Modified Eagle Medium (DMEM), 10% FBS, 1% Penicillin/Streptomycin and 0.2% Collagenase for 70 minutes followed by 0.25% dispase for 10 minutes.^[55]^ hFAPs were isolated using FACS Aria Fusion with SYTOX™ Blue Dead Cell Stain (ThermoFisher, #S34857) and further isolated using CD31-/CD45-/CD56-/CD29-/CD34+ markers.^[56]^ hFAPs were then cultured in Ham’s F-10 Nutrient Mix (F-10) plus 10% FBS, 10 µg/mL basic fibroblast growth factor, and 1% Penicillin/Streptomycin until encapsulation.

### Encapsulation of hFAP Cells into HyA Hydrogels

hFAP cells were trypsinized from 2D culture and encapsulated at a density of 3 million cells/mL in HyA hydrogels. hFAP cells and crosslinker solution were mixed simultaneously into the macromer precursor solution and injected into 12 mm Millicell cell culture inserts with a 0.4 µm pore size (Millipore Sigma, #PIHP01250). Cell-gel constructs were allowed to crosslink completely for 1 hour in a 37°C, humidity-controlled incubator, after which cell culture media was added to each well.

### Cell culture, viability, and proliferation

Cell-gel constructs received daily media changes. To assess cell viability, samples were washed twice with PBS and incubated for 1 hour in media supplemented with 2 µM calcein AM and 4 µM ethidium homodimer at the desired time points. Samples were washed in PBS twice and imaged using a fluorescent Lionheart microscope (Agilent BioTek, Santa Clara, CA). hFAP proliferation was assessed after 0 (immediately after encapsulation), 1, 4, 7, and 10 days in culture using the Alamar Blue assay per the manufacturer’s instructions. Briefly, cell-gel constructs were incubated with 10% Alamar Blue in cell culture media for 2 hours, after which media from each well was collected and its fluorescence measured to assess relative metabolic activity.

### Immunocytochemistry and Quantification

Hydrogel-cell constructs were fixed at the desired time points using 4% (v/v) paraformaldehyde for 30 min, washed with PBS, and permeabilized with 0.05% PBST for 1 hour. After blocking with 5% BSA for 1 hour, samples were incubated overnight at 4°C with a 1:500 dilution of primary antibody for UCP1 (Abcam, ab155117) and mouse anti-αSMA (Abcam, ab7817) in 5% BSA. After washing the samples with PBST, samples were incubated with DAPI (1:1000), rhodamine phalloidin (1:400, Invitrogen, R415), and secondary antibodies for UCP1 and αSMA (1:1000, Invitrogen, A21245 and A11001 respectively) overnight at 4°C. Prior to imaging, cell-gel constructs were washed with PBST and imaged using a PerkinElmer (Waltham, MA) Opera Phenix confocal microscope equipped with a 20X water objective. Cell morphological and fluorescent intensity properties were assessed by creating 3D projections of 1100 µm z-stacks produced at each imaging region (x-y location, n ≥ 20) within the hydrogels. Encapsulated hFAPs were identified as singular objects within each *z*-stack using the DAPI channel to identify the nucleus of each cell and phalloidin to define the boundaries of the surrounding cytosol. These object boundaries were used to calculate morphological properties such as cell volume and to define regions within which to determine cell UCP1 and αSMA intensity values (**SI Figure 2**).

### Human angiogenesis protein profiler array

Encapsulated hFAPs were evaluated for their expression of a variety of angiogenic proteins using a Proteome Profiler Human Angiogenesis Array Kit (R&D Systems, #ARY007) at 1-, 8-, and 14-days post-encapsulation. Briefly, HyA hydrogels containing encapsulated hFAPs were degraded in 3000 U/mL hyaluronidase from bovine testes (Sigma, #37326-33-3) to release the entrapped cells. Cells were purified from the degraded HyA/cell mixture before lysing the cells and isolating proteins using RIPA lysis and extraction buffer (ThermoFisher, #89900 and #89901) with added protease and phosphatase inhibitors (ThermoFisher, #PIA32965). Protein solutions were then added to each array, following the manufacturer’s instructions. Arrays were visualized using an Analytik Jena UVP ChemStudio Series Imager (Jena, Germany). Relative protein expression was quantified via measurement of pixel intensity after subtracting background signal followed by comparison of pixel intensities between corresponding time points for each protein in ImageJ.

### hFAP secretion of IL-10

A Quantikine solid phase sandwich ELISA kit for IL-10 (R&D Systems, #D1000B) was performed on the media collected from samples at 1- and 7-days post-encapsulation following the manufacturer’s instructions to quantify the relative amount of IL-10 in each sample group.

### Surgical Approach

All animal procedures were approved by the San Francisco VA Medical Center Institutional Animal Care and Use Committee (Kim 24-006) and performed under IRB protocol. Ten male 9-month-old NSG mice underwent unilateral right suprascapular nerve denervation (TTDN) and combined supraspinatus (SS) and infraspinatus (IS) tendon transection, described previously.^[57–59]^ The SS was tagged with 7.0 suture for later identification for repair. After 6 weeks of observation, each mouse received a delayed partial rotator cuff repair under isoflurane anesthesia by reconnecting the SS with 7.0 suture to the greater tubercle of the humerus through a single bone tunnel created with a 26-gauge needle. The mice were then separated and injected intramuscularly with either 10µl saline (PBS, n=5) or 10µl HyA-hydrogel (n=5). The hydrogel was crosslinked immediately prior to injection by mixing sterile macromer and crosslinker precursor solutions. The mixture (or PBS as a control) was then intramuscularly injected at five distinct locations along the length of the supraspinatus using a syringe with a 23-gauge needle. Animal wounds were closed and closely monitored for post-surgical survival. Mice were humanely sacrificed at 6 weeks post-repair and bilateral SS muscles were harvested, weighed for percent muscle weight loss, and immunofluorescently analyzed to quantify fibrosis, fat index, muscle fiber cross-sectional area (CSA), and vascularity.

### Muscle Harvesting and Histology

Mice were humanely sacrificed in CO_2_ chamber and the supraspinatus and infraspinatus rotator cuff muscles were harvested bilaterally. To ensure that the same portion of the muscle was analyzed between groups, all histological sections were taken from the central region of the muscle belly, and all muscle samples were frozen and mounted in the same orientation. The samples were prepared on a cork, flash frozen in isopentane, sectioned to 7 µm using a cryostat, and stored at -80°C. For immunofluorescence, sections were fixed with 4% paraformaldehyde for 10 minutes, washed in PBST (1x PBS with 0.1% TritonX-100), blocked for 1 hour in 5% BSA, and incubated with rabbit anti-laminin antibody (Sigma, L9393) and anti-CD31 antibody (ThermoFisher, 48-0319-42) at 4°C overnight. Slides were then washed in PBS and incubated with goat anti-rabbit secondary antibodies (Abcam, ab150115) for 1 hour at room temperature. The tissue sections were stained with DAPI and then mounted with Fluoromount G. Oil Red O staining was completed to show the amount of FI by the percentage of area stained for fat per field of view. Masson’s trichrome staining indicated degree of fibrosis by the percentage of collagen staining fibers per field of view.

### Statistical Analysis

Data are presented as mean ± standard deviation of replicates. Frequency sweep (n=7) and stress relaxation (n=4) rheological data of HyA hydrogels are presented as average values with error bars representing the standard deviation between replicates. Statistical analysis of hydrogel parameters after swelling in different solvents (n=3) was completed using a one-way ANOVA with Tukey’s multiple comparisons test. Analysis of *in vitro* immunostaining quantification (n=25) was conducted via a one-way ANOVA with Dunnett’s multiple comparisons test. *In vitro* IL-10 secretion data at day 1 (n=5) vs. day 7 (n=3), *in vivo* paired muscle weight loss, and *in vivo* CSA loss data were compared using a Student’s t-test. A two-way ANOVA with Tukey’s multiple comparisons test was used for statistical analysis between groups for *in vivo* experiments (n=5 animals per treatment, n=3 histological sections per animal). Prism 10 software (version 10.2.0, GraphPad Software, Inc.) was used for all statistical testing and figure creation except for the proteome profiler relative protein expression data, which was plotted and visualized in Microsoft Excel. Statistical significance was defined by *α* < 0.05.

## Acknowledgements

This work was supported in part by the California Institute for Regenerative Medicine (CIRM DISC2-13201), the NIH National Institute of Arthritis and Musculoskeletal and Skin Diseases (1R01AR072669), and the VA ORD CSR&D Merit Review grant (I01 CX002200). For ^1^H NMR material characterization, we thank Drs. Hasan Celik, Raynald Giovine, and Pines Magnetic Resonance Center’s Core NMR Facility (PMRC Core) for spectroscopic assistance. The instrument used in this work is supported by the National Science Foundation under Grant No. 2018784. This material is based upon work supported by the National Science Foundation Graduate Research Fellowship Program under Grant No. DGE 2146752. Any opinions, findings, and conclusions or recommendations expressed in this material are those of the author(s) and do not necessarily reflect the views of the National Science Foundation. Some images were created with BioRender.

## Author Contributions

K.E.H., B.T.F., and X.L. conceived and designed the research and experiments, supervised the project development, and funded the study. M.P. completed the material characterization. M.P., A.K., and D.N. performed *in vitro* studies using hFAPs isolated by S.G. and A.W from muscle tissues. M.D. and M.L. performed the *in vivo* surgeries and hydrogel injections, and M.D., A.N., and P.N. completed the histological staining. M.P., A.W., B.T.F., X.L., and K.E.H. wrote the manuscript with discussions and improvements from other authors.

## Supporting Information

**SI Figure 1:**
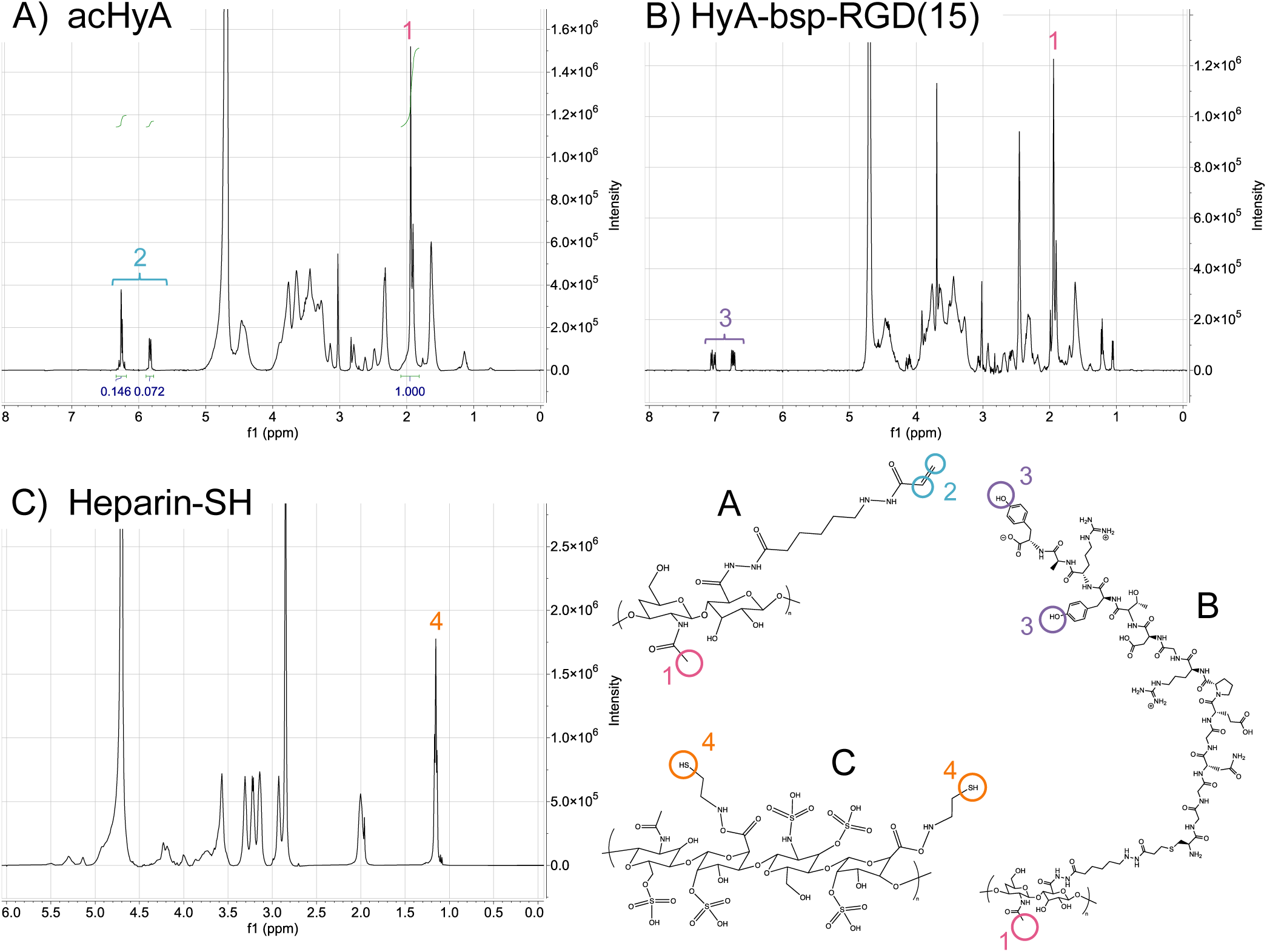
NMR Spectra of Hydrogel Components. A) acHyA. B) acHyA conjugated to bsp-RGD(15). C) Thiolated heparin.

**SI Figure 2:**
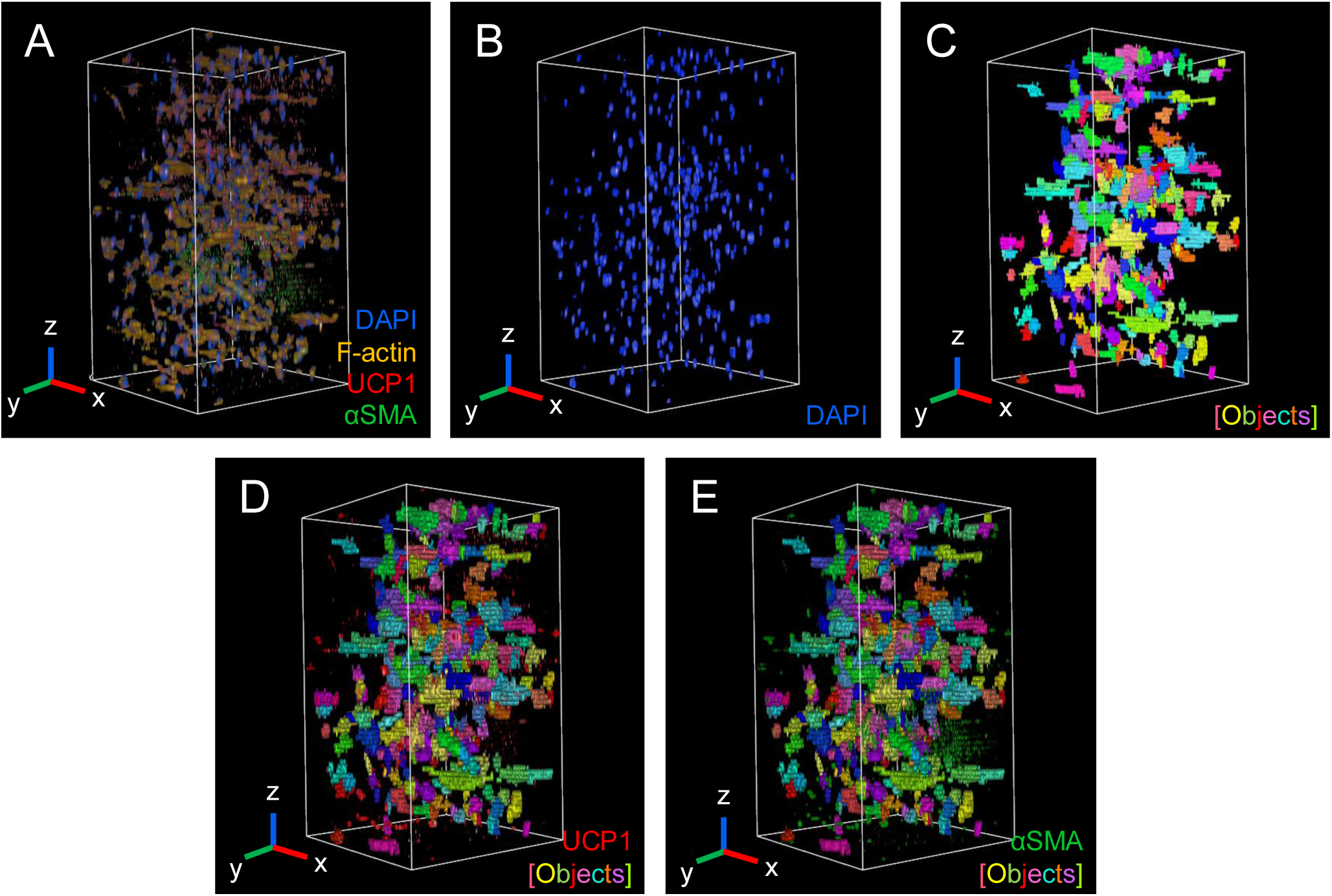
3D Cell-Gel Analysis Pipeline. 3D projection, cell detection, and analysis functions were utilized to quantify hFAP behavior inside HyA hydrogels over time. A) 3D projection of a representative 1,100 µm z-stack produced at each imaging region within a hydrogel, consisting of 4 channels: DAPI (blue, nucleus), rhodamine (yellow, F-actin), Alexa 488 (green, αSMA), and Alexa 647 (red, UCP1). B) The DAPI channel was used for the identification of nuclei, each of which became an independent object for further segmentation (all objects highlighted in blue). C) The rhodamine (F-actin) channel was used to identify the surrounding cytosol for each object identified in B (cell boundaries highlighted in assorted colors). These boundaries were then used to quantify morphological properties (cell volume, eccentricity) or fluorescent intensity for D) UCP1 and E) αSMA within each cell.

